# Aggregation of the Amyloid-β Peptide (Aβ40) within Condensates Generated through Liquid-Liquid Phase Separation

**DOI:** 10.1101/2023.12.23.573169

**Authors:** Owen M. Morris, Zenon Toprakcioglu, Alexander Röntgen, Mariana Cali, Tuomas P. J. Knowles, Michele Vendruscolo

## Abstract

The deposition of the Aβ peptide into amyloid fibrils is characteristic of Alzheimer’s disease. As it has been recently observed that the process of amyloid aggregation can take place within an intermediate liquid-like condensed phase, we investigated whether Aβ could undergo liquid-liquid phase separation, and whether Aβ amyloid aggregation could take place within Aβ liquid condensates. By using a microfluidic protocol, we observed that the 40-residue form of Aβ (Aβ40) can undergo liquid-liquid phase separation, and that accessing a liquid intermediate state enhances primary nucleation and enables Aβ40 to readily self-assemble into amyloid fibrils. These results prompt further studies to investigate the possible role of Aβ condensates in the aggregation of this peptide in Alzheimer’s disease.

## Introduction

Alzheimer’s disease is a progressive neurodegenerative disorder, accounting for approximately two-thirds of all cases of dementia worldwide, with an estimated yearly societal cost exceeding $1 trillion (1, 2). The aggregation of the amyloid-β peptide (Aβ) into amyloid fibrils and amyloid plaques has long been associated with Alzheimer’s disease, although the exact nature of the association has not yet been fully clarified (3-5). Aβ is an intrinsically disordered protein fragment that originates from the proteolytic cleavage of the amyloid precursor protein (APP) (3, 4). A reaction by which Aβ self-assembles from its monomeric soluble form into amyloid aggregates has been extensively studied *in vitro* (5-7). Kinetic analysis has revealed that Aβ monomers initially bind to one another, in a process known as primary nucleation, to form oligomeric assemblies. These oligomeric assemblies then grow into amyloid fibrils through the addition of further Aβ monomers to their ends, in a process known as elongation. The amyloid fibrils, in turn, act as catalytic surfaces for the formation of new oligomeric assemblies. This surface-catalysed process is known as secondary nucleation, and it is the dominant mechanism for the proliferation of Aβ aggregates (5, 6, 8).

Recently, however, an alternative pathway to amyloid formation has emerged, sometimes referred to as the condensation pathway (9), whereby a protein aggregates into amyloid fibrils by initially undergoing a liquid-liquid phase separation process (8). The liquid-liquid phase separation of proteins involves the separation of a homogeneous liquid phase into two coexisting liquid phases of different densities (10-14). Within the high-density phase, nucleation processes, which are highly concentration dependent, are greatly accelerated, resulting in the rapid formation of amyloid aggregates. This condensation pathway to aggregation has been reported for proteins involved in neurodegenerative diseases, including α-synuclein, tau and FUS (15-20). These observations, together with the suggestion that a large fraction of human proteins may undergo phase separation (21), and the growing number of proteins for which this process has been reported *in vivo* (22), are prompting the question of whether Aβ could also undergo amyloid aggregation from within liquid condensates.

In this study, we used an *in vitro* microfluidic assay combined with fluorescence microscopy to observe the liquid-liquid phase separation of Aβ40 in the presence of claramine, a small molecule in the aminosterol class (23, 24) that has recently been reported to facilitate the liquid-liquid phase separation of α-synuclein (25), and the subsequent formation of Aβ40 amyloid fibrils within the dense liquid phase.

## Results and Discussion

### Liquid-liquid phase separation of Aβ40

A general workflow for investigating the liquid-liquid phase separation of Aβ40 by using microfluidics is outlined in **Figure 1**. With this workflow, we found that incubating Aβ40 with claramine results in liquid-liquid phase separation. Claramine, which is a synthetic analogue of the natural aminosterol trodusquemine, is a small cationic molecule consisting of a polyamine (spermine) moiety covalently bound to a cholestane sterol group. Aminosterols were first discovered when squalamine was isolated from stomach and liver extracts from *Squalus acanthias* and shown to have antimicrobial properties (26, 27). Recently, there has been a growing interest in the use of aminosterols as potential therapeutics in neurodegenerative diseases (28, 29). This interest has stemmed from studies reporting the ability of squalamine and related compounds to reduce the cytotoxicity of α-synuclein and Aβ oligomers (24, 30-34).

**Figure 1.**
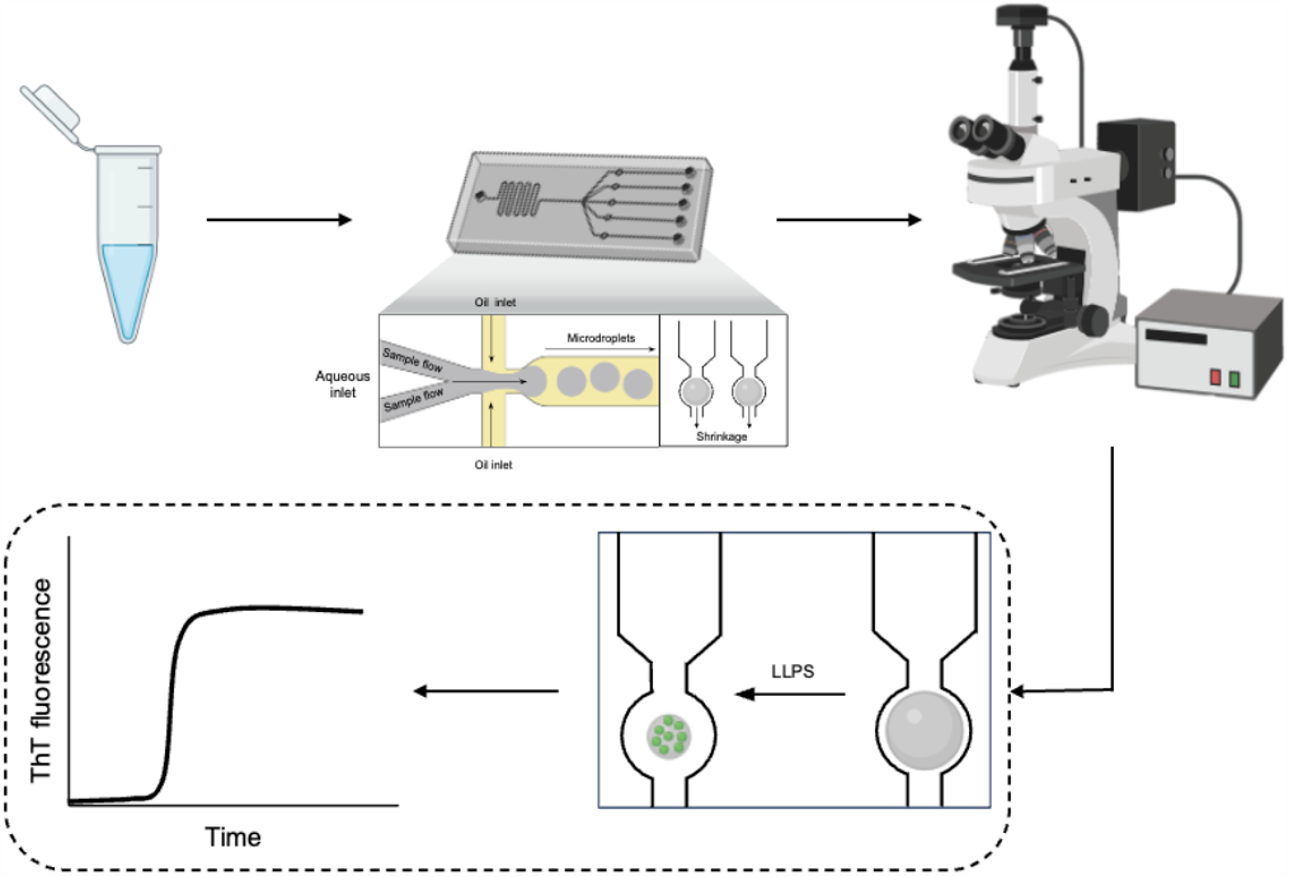
Schematic of microfluidic assay to monitor liquid-liquid phase separation and amyloid aggregation of Aβ40. An aqueous Aβ40 sample (3-5 μl) is initially injected into the microfluidic device. Aqueous microdroplets are subsequently generated and entrapped within the device (35). Fluorescence microscopy is then utilised to monitor the shrinkage of these microdroplets over time, and to observe the liquid-liquid phase separation of Aβ40. time-lapse images of the microdroplets may be analysed using ImageJ (42), and the liquid-liquid phase separation and aggregation of Aβ40 can be characterised. Depiction of the microfluidics device and fluorescence microscope have been adapted from BioRender.

Aβ40 samples with varying amounts of claramine were injected into a microfluidic device (see Methods) to investigate the propensity of Aβ40 to undergo liquid-liquid phase separation (35). This approach involves the use of an array of microfluidic traps whereby aqueous sample-containing microdroplets are confined and studied over time (**Figure 1**). To avoid microdroplet coalescence and surface effects, a fluorinated oil was used (16). Fluorescence microscopy was utilised to monitor the time-dependent shrinkage of these microdroplets, and to observe Aβ40-containing condensates generated by liquid-liquid phase separation. The liquid-liquid phase separation of Aβ40 was observed within the microdroplets for various stoichiometries of Aβ40:claramine (**Figure 2A**), using a small proportion of AF488-labeled Aβ40 (see Methods). These individual microdroplets containing Aβ40 condensates were also visualised using confocal microscopy (**Supplementary Figure 1**). A 1:1 ratio of Aβ40:claramine was found to be most effective for promoting liquid-liquid phase separation. This finding was confirmed by calculating the saturation concentration (C_sat_), which is the concentration of Aβ40 required to generate biomolecular condensates (8) (**Figure 2B**).

**Figure 2.**
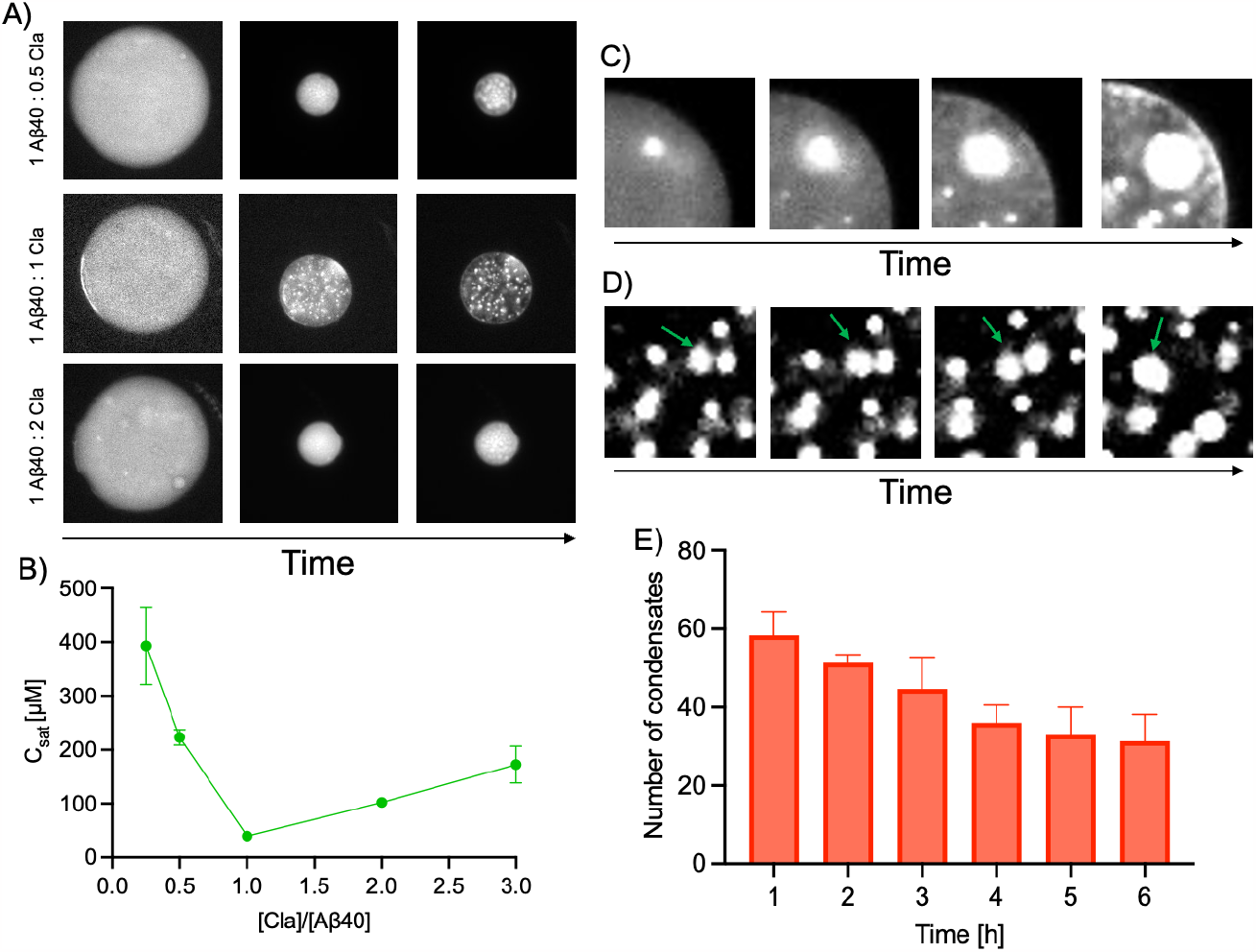
Formation of Aβ40 condensates by liquid-liquid phase separation. **(A)** Time-lapse fluorescence microscopy images (excitation at 490 nm) of microdroplets displaying Aβ40 condensates as the individual microdroplets shrink. **(B)** Concentration at which liquid-liquid phase separation was observed (C_sat_) for Aβ40 incubated with various concentrations of claramine. Three data points were averaged for each C_sat_ value. **(C)** Fluorescence microscopy time-lapse images displaying Ostwald ripening of Aβ40 condensates. **(D)** Fluorescence microscopy time-lapse images displaying coalescence events occurring between Aβ40 condensates. **(E)** Bar chart showing the decrease in the total number of condensates over time for a 1:1 ratio of Aβ40 to claramine. Analysed condensates were selected based on those being in-focus within the fluorescence microscopy images. Data are shown as mean ± SD of n=3.

Monitoring the Aβ40 condensates indicated typical liquid-like behaviour. Aβ40 condensates were observed to undergo Ostwald ripening, where larger coacervates grow rapidly at the expense of smaller coacervates (**Figure 2C**). Furthermore, coalescence of Aβ40 condensates was also observed, as condensates of increasingly larger sizes were produced (**Figure 2D** and **Supplementary Figure 2**). Both processes can be observed, along with the maturation of Aβ40 condensates when monitoring the microdroplets within the microfluidic device over time (**Supplementary Videos 1 and 2**). Furthermore, a liquid-like behaviour is corroborated by observing the number of Aβ40 condensates within an individual microdroplet. As the microdroplet shrinks and liquid-liquid phase separation is initiated, a decrease in the total number of condensates is observed due to coalescence events (**Figure 2E**). Eventually, the number of Aβ40 condensates reaches a plateau, as the individual condensates mature and lose their liquid properties as they become more gel-like and exhibit solid-like behaviour.

### Aggregation of Aβ40 within liquid condensates

Next, we sought to investigate the propensity of Aβ40 to aggregate within the liquid condensates. To study the self-assembly of Aβ40 into the amyloid state, we added the amyloid binding dye Thioflavin T (ThT) at 20 μM. Once ThT is bound to the β-sheet core of amyloid fibrils, it becomes fluorescent at 490 nm (36, 37). Additionally, the use of labelled Aβ40 was substituted for wild-type Aβ40 to eliminate any crosstalk between the emission intensity of AF488-Aβ40 and the emission intensity of ThT.

Microfluidic experiments involving 7 μM of Aβ40 in conjunction with a 0.25-3 times stoichiometry of claramine were conducted in the presence of ThT. As the individual microdroplets within the microfluidic device shrank with time, liquid-liquid phase separation was once again observed between Aβ40 and all stoichiometries of claramine (**Figure 3A**). Additionally, Aβ40 subsequently underwent self-assembly within its condensed liquid state with all ratios of Aβ40:claramine, as indicated by an increase in the ThT intensity within Aβ40 condensates. By monitoring this process with fluorescence microscopy, we estimated the critical concentration of Aβ40 within each microdroplet required to initiate the self-assembly of Aβ40 within a condensate (**Figure 3B**). Similar to the findings described above, a 1:1 Aβ40:claramine ratio was optimal for promoting Aβ amyloid formation within the liquid condensates. At lower claramine stoichiometries, we observed that a sharp increase in the critical concentration of Aβ40 was required to initiate the self-assembly, whereas at stoichiometries beyond the 1:1 condition, a gradual increase in Aβ40 critical concentration was needed (**Figure 3B**).

**Figure 3.**
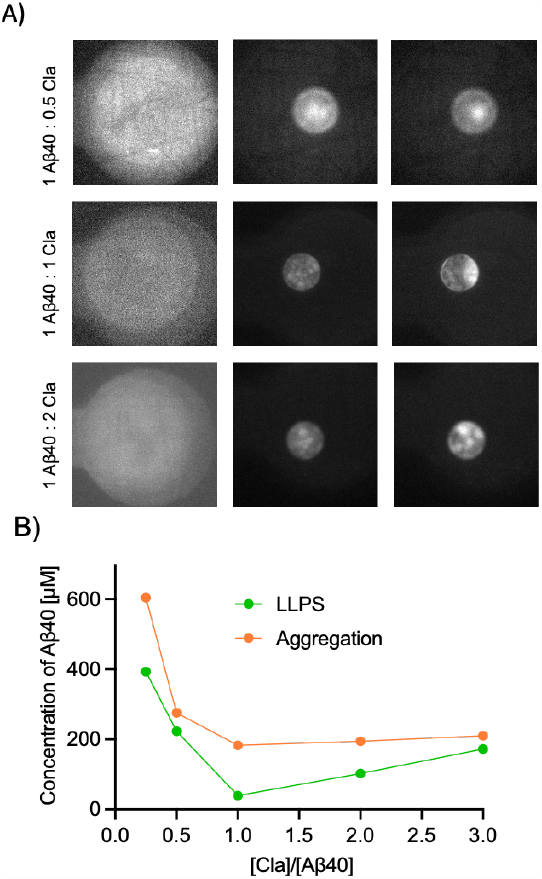
Amyloid aggregation of Aβ40 within liquid condensates. **(A)** Time-lapse fluorescence microscopy images (excitation at 490 nm) of Aβ40 undergoing aggregation within condensates. **(B)** Phase diagram of Aβ40 and claramine. The green line displays the concentration at which Aβ40 undergoes liquid-liquid phase separation at various Aβ40:claramine stoichiometries. The orange line displays the concentration of Aβ40 at which the initial stages of aggregation are observed within the condensed liquid phase. Data are shown as mean of n=3.

Combining the plots for the C_sat_ of Aβ40 (**Figure 2B**) and its critical concentration to initiate its self-assembly (**Supplementary Figure 3**) generates a phase boundary of Aβ40 in the presence of claramine (**Figure 3B**). This boundary details the concentrations of Aβ40 required to induce its liquid-liquid phase separation and form coacervates at each stoichiometry, along with the Aβ40 concentration required to initiate aggregation within the coacervate.

### Kinetic analysis of Aβ40 aggregation within liquid condensates

Further analysis of these microfluidic experiments involving Aβ40 and ThT yielded a series of kinetic aggregation curves for each Aβ40:claramine ratio. These individual kinetic traces follow a characteristic sigmoidal pattern, whereby a rapid exponential growth phase is preceded by a lag phase and followed by a plateau phase (5, 8). The raw kinetic traces for each Aβ40:claramine ratio were then normalised (**Figure 4C**). The normalised kinetic curves support the conclusion that a 1:1 ratio of Aβ40 to claramine aggregates within the condensed liquid state more readily than other stoichiometries. This finding was also reinforced by extracting the half-time of aggregation, which is the time taken for the fibrillar mass to reach half of its plateau value (**Figure 4B**).

**Figure 4.**
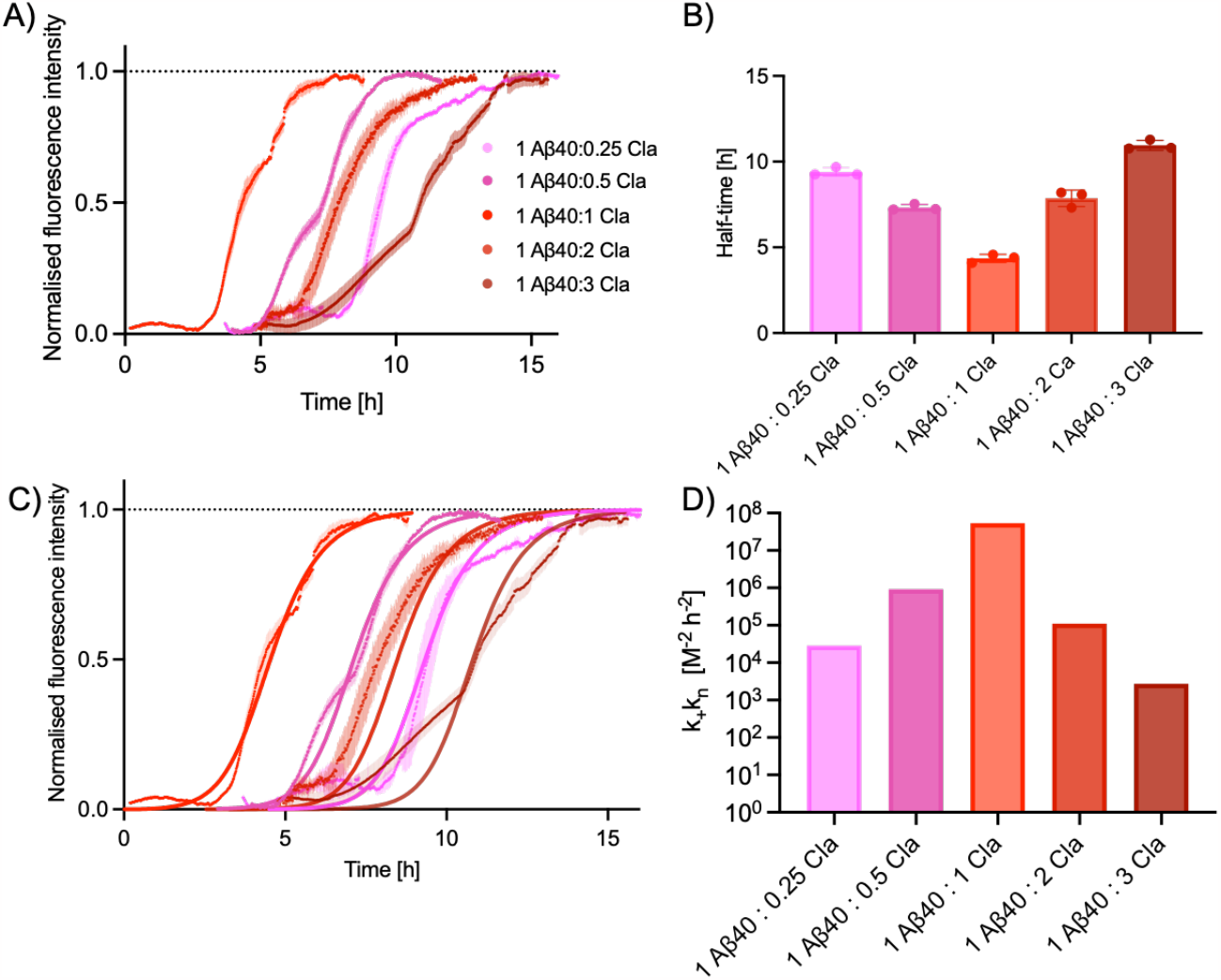
Kinetics of amyloid aggregation of Aβ40 within liquid condensates. **(A)** Normalised kinetic traces of Aβ40 aggregation within liquid condensates at various Aβ40:claramine stoichiometries. The legend indicates the colour code used throughout the figure for the Aβ40:claramine stoichiometry. **(B)** Bar chart of the half-time of Aβ40 aggregation within liquid condensates for various Aβ40:claramine stoichiometries. **(C)** Normalised kinetic traces for Aβ40 aggregation within liquid condensates at various Aβ40:claramine stoichiometries. The solid lines represent the fits to the kinetic data, which was performed using AmyloFit (38). Data are shown as mean ± SD of n=3 (A–C). **(D)** Bar chart of the value of the rate parameter for primary nucleation, k_+_k_n_, as a function of the Aβ40:claramine stoichiometry.

To elucidate the microscopic processes that dominate the macroscopic aggregation behaviour, we used AmyloFit, a global fitting platform that compares the integrated rate laws to experimental kinetic data (38). By employing a chemical kinetics framework, we were thus able to elucidate the rate constants of the microscopic steps involved in this aggregation process through the condensation pathway (16). These microscopic steps include primary nucleation (k_n_), elongation (k_+_) surface-catalysed secondary nucleation (k_2_). The reaction order of the primary process is denoted by (n_c_), while the reaction order of the secondary process in the pathway is denoted by (n_2_). Importantly, only two key rate parameters control the time course of aggregation. These are the combinations of rate constants k_+_k_n_ and k_+_k_2_, which describe aggregate proliferation through primary and secondary nucleation, respectively (39).

We first obtained the rate parameters for the 1:1 Aβ40:claramine ratio, and then the rate parameter for secondary nucleation, k_+_k_2_, was kept constant throughout the analysis at 3.61x10^15^ M^-3^h^-2^. The rate parameter for primary nucleation, k_+_k_n_, for each system was then obtained (**Figure 4D**). We found that small differences in the Aβ40:claramine ratio can greatly modulate the rate parameter for primary nucleation, in some cases changing it by up to 5 orders of magnitude. Furthermore, a non-monotonic behaviour between the nucleation rate and the Aβ40:claramine ratio was observed, with the maximum rate being obtained when the Aβ40:claramine ratio was 1. Our data suggest the presence of two regimes. In the low Aβ40:claramine ratio, the primary nucleation rate is supressed (**Figure 4D**). This implies that a critical amount of claramine is needed to induce condensate formation and subsequent aggregation within the condensates. In the high Aβ40:claramine ratio regime, the primary nucleation rate is again supressed. This suggests that although claramine can induce phase separation and aggregation, if it is in excess, it can also interfere with these processes. We note that claramine has already been shown to stabilise the condensed phase and affect the primary nucleation rate of another amyloidogenic protein, α-synuclein (25).

To identify the amyloid aggregates formed within liquid condensates, transmission electron microscopy (TEM) was utilised to confirm whether Aβ40 formed fibrils for all stoichiometric ratios. Amyloid fibrils were observed for all conditions (**Figure 5**).

**Figure 5.**
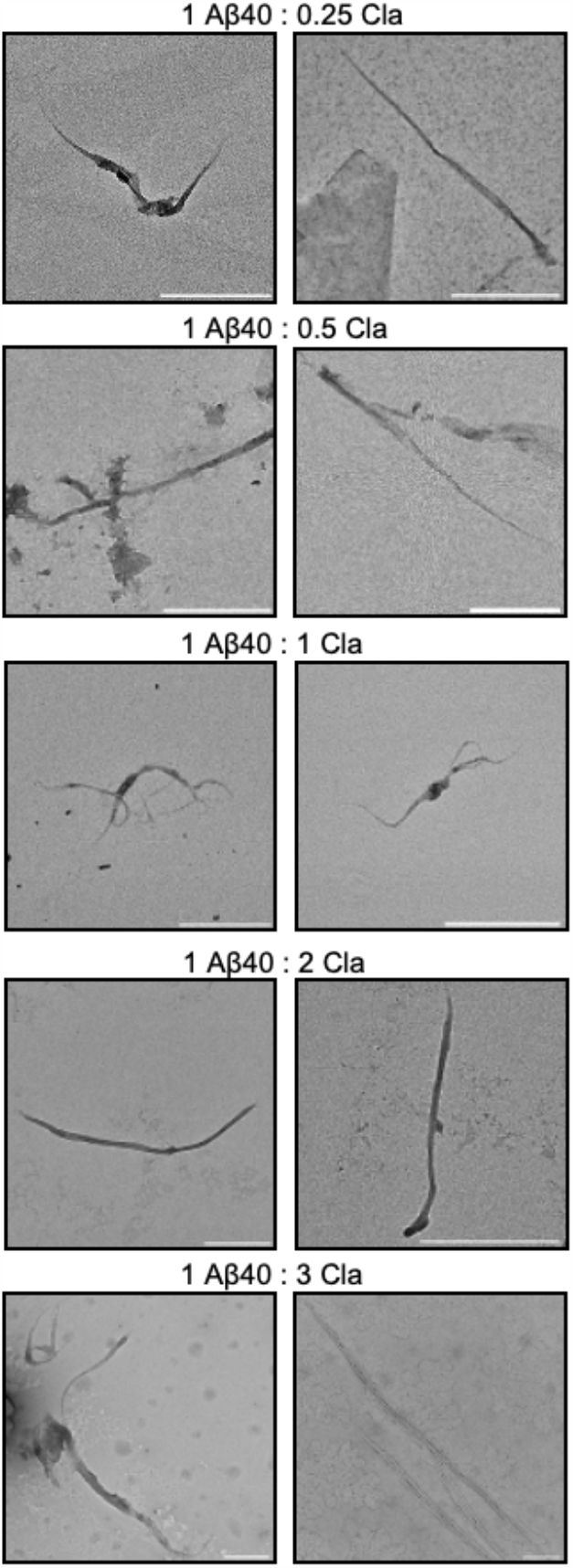
Transmission electron microscopy (TEM) images of Aβ40 fibrils. Fibrillar aggregates of Aβ40 were imaged in the presence of various stoichiometries of claramine. Two micrographs were displayed for each condition. Scale bar shown in all images is 1 μm.

## Conclusions

We have reported that Aβ40 can form amyloid aggregates through the condensation pathway, as summarised in **Figure 6**. In this pathway, Aβ40 first populates a condensed state by undergoing liquid-liquid phase separation. The resulting condensates then readily convert into amyloid fibrils because the high concentration of protein within the condensates greatly enhances primary nucleation. For comparison, the condensation pathway differs from the well-established deposition pathway since in the latter the intermediate states are disordered oligomeric assemblies (**Figure 6**). We anticipate that the condensation pathway of Aβ will be further investigated due to its possible relevance in amyloid formation *in vivo*, and ultimately in Alzheimer’s disease.

**Figure 6.**
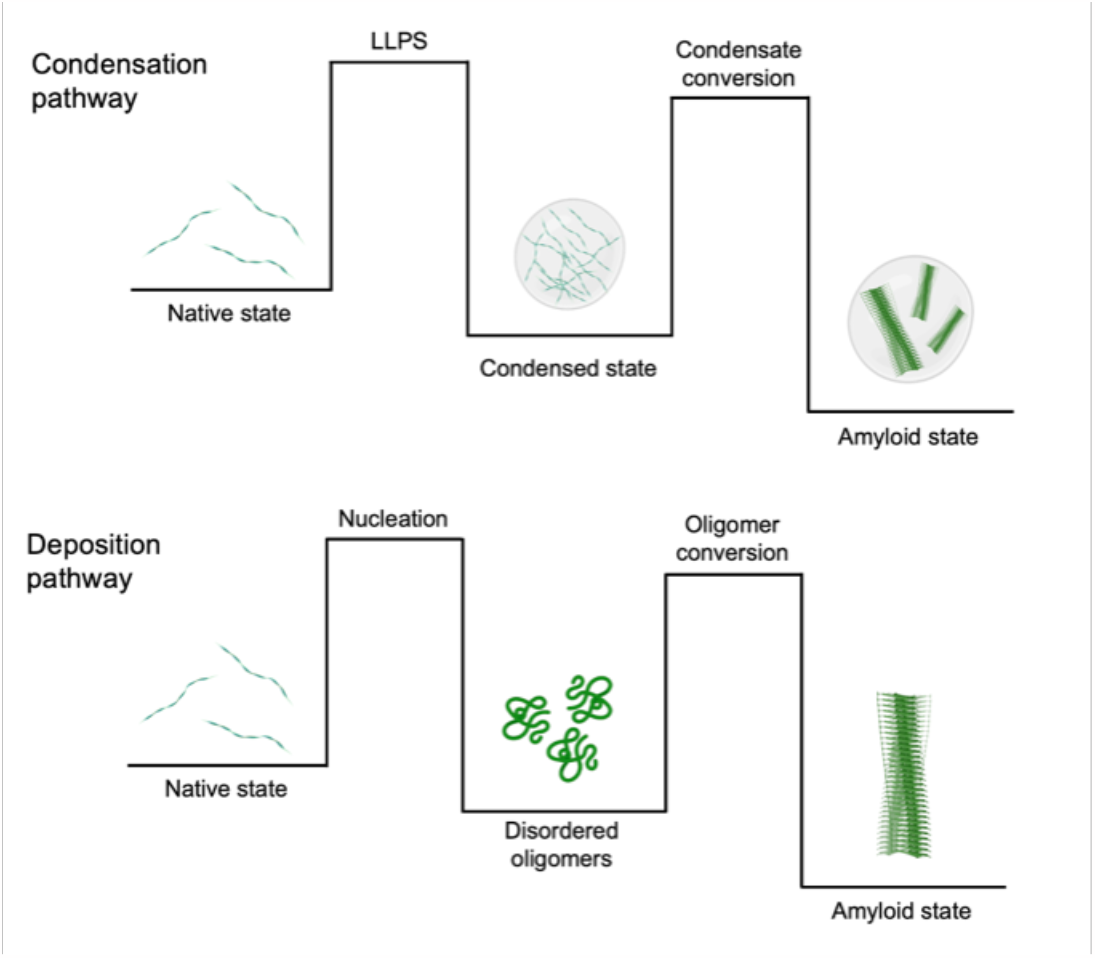
Schematic comparison of the condensation and deposition pathways of aggregation of Aβ40. The conversion between the native and amyloid states along the condensation pathway takes place through an intermediate condensed state. Along the deposition pathway, the conversion takes place through the formation of disordered oligomers.

## Materials and Methods

### Purification and fluorescent labelling of Aβ40

Human wild-type Aβ40 and its cysteine-bearing variant (insertion at position 2) were expressed in *E. coli* BL21 Gold (DE3) competent cells, transformed with a pT7 plasmid encoding the relevant Aβ40 variant. *E. coli* cells were induced at an optical density (OD) of 0.8 using 1 mM isopropyl β-D-1-thiogalactopyranoside (IPTG). The Aβ40 variants were subsequently purified in 50 mM Tris-HCl pH 7.4 using a previously established protocol to achieve consistent properties (40, 41). All Aβ40 was aliquoted, flash-frozen in liquid nitrogen, lyophilised, and stored at -80 °C. Lyophilised Aβ40 was redissolved in 50 mM Tris-HCl pH 7.4 on ice at a stock concentration of (50 μM) before each experiment.

The cysteine-bearing Aβ40 variant was redissolved in 50 mM NaPi pH 7.5 and labelled with an 8-fold molar excess of Alexa Fluor™ 488 C_5_ Maleimide (AF488; Invitrogen Life Technologies) and incubated at room temperature for 2 h (**Supplementary Figure 3**). Labelled Aβ40 was separated from unbound dye and Aβ40 dimers by size exclusion chromatography (23 ml Superdex 75 10/300 GL column) using a flow rate of 0.7 ml/min 50 mM Tris-HCl pH 7.0. The concentration of labelled Aβ40 was determined from the Beer-Lambert law by measuring the UV-Vis absorbance at 495 nm using an extinction coefficient ε= 73,000 M^-1^ cm^-1^ for AF488.

### Microfluidic liquid-liquid phase separation assay

#### Liquid-liquid phase separation assay composition

7 μM of Aβ40 (10% AF488-labelled) was combined with varying stoichiometries of claramine (Sigma-Aldrich) and 5% polyethylene glycol (PEG) (50% w/w 10 kDa, Thermo Fisher Scientific) in 50 mM Tris-HCl pH 7.4. Aqueous microdroplets consisting of a varying mixture of PEG/claramine and peptide were generated and confined in the microfluidic trapping device for subsequent imaging (35).

#### Fabrication of microfluidic devices

The fabrication process of the microfluidics was taken from a previously established protocol (35). A soft photolithographic process was used to fabricate the master through which microfluidic devices were made. A 50 μm photoresist (SU-8, 3050, MicroChem) was spin-coated onto a silicon wafer. This was soft baked at 95 °C for 3 min. A film mask was placed on the wafer and system was exposed to UV light to induce polymerization. The wafer was then baked at 95 °C for 30 min. Finally, the master was placed into a solution of propylene glycol methyl ether acetate (PGMEA, Sigma-Aldrich), which helped in the development process. Elastomer polydimethylsiloxane (PDMS) with curing agent (Sylgard 184, DowCorning, Midland, MI) was mixed at a ratio of 10:1 to fabricate he devices. This mixture was then incubated at 65 °C and cured for a total of 3 h. Once hardened, the PDMS was peeled off the master, and holes were punched into the PDMS, which acted as inlets and outlets. Finally, the PDMS slab was bound to a glass slide by treatment with a plasma bonder (Diener Electronic, Ebhausen, Germany).

#### Formation and confinement of microdroplets

Syringe pumps (neMESYS, Cetoni, Korbussen, Germany) were used to control the flow rates within the microchannels. The Aβ40 solution was introduced to the device at the first junction. At the second junction, the oil phase, which consisted of fluorinated oil (Fluorinert FC-40, Sigma-Aldrich) and 2% w/w fluorosurfactant (RAN biotechnologies) intersected the aqueous phase, which resulted in water-in-oil microdroplets being formed. Following microdroplet generation, microdroplets were entrapped within the microfluidic device. Briefly, microdroplets were directed towards an array of traps whereby once a microdroplet is driven within the microfluidic confinement, it is unable to escape unless a pressure is applied from the outlet. Microdroplets were then incubated at room temperature to allow for shrinkage. This resulted in an increase of the local concentration of Aβ40 and PEG within the microdroplets which led to Aβ40 phase separation (**Supplementary Figures 4 and 5**). The water-in-oil microdroplets and the Aβ40 phase separation was monitored using fluorescence microscopy (Bright-field and GFP-channel, 488 nm).

#### Calculation of Aβ40 concentration during phase separation

The concentration of Aβ40 required to induce liquid-liquid phase separation was obtained by calculating the ratio of the microdroplet volume following trapping and at the point of phase separation, i.e. the point at which condensates start appearing within the water-in-oil microdroplet (16). By multiplying the initial monomeric Aβ40 concertation by the value of this ratio, the Aβ40 concentration at the point of phase separation could be determined.

### Aβ40 aggregation assay within liquid condensates

7 μM Aβ40 was incubated with varying stoichiometries of claramine and 5% PEG (50% w/w 10 kDa, Thermo Fisher Scientific) in the presence of 20 mM ThT (Sigma-Aldrich) in 50 mM Tris-HCl pH 7.4. Approximately 5 μl of the assay mixture was injected into the microfluidics device, where microdroplets were entrapped and imaged at 490 nm using fluorescence microscopy. Images were processed using ImageJ (NIH), where an individual microdroplet is analysed.

### Confocal microscopy

Approximately 5 μl of Aβ40 sample was injected into the microfluidic device using the method outlined in the *Formation and confinement of microdroplets* section and imaged using a Leica TCS SP8 inverted confocal microscope using a 20x objective (Leica microsystems). The excitation wavelength was set at 490 nm. Images were processed using ImageJ (NIH).

### Transmission electron microscopy (TEM)

TEM was performed at the Cambridge Advanced Imaging Centre using a Talos F200X G2 transmission electron microscope operating at 80-200 kV. TEM images were acquired using a Ceta 16M CMOS camera. TEM grids (continuous carbon film on 300 mesh Copper grid) were glow discharged using Quorim Technologies GloQube instrument at a current of 25 mA for 60 s. Samples were negatively stained using a 2% w/v uranyl acetate solution for 40 s and air-dried before imaging.

### AmyloFit kinetic analysis

All kinetic data obtained were analysed using AmyloFit (38). All datasets were initially normalised prior to upload. The half-times were calculated from the time at which the fibrillar mass reached half its maximum value, i.e., the time at which the normalised fluorescence intensity reached 0.5. Each experiment was repeated three times and averaged before fitting. From the data, a secondary nucleation dominated model was found to be the best model to use. The fitting was initially performed on the 1:1 Aβ40:claramine ratio and subsequently, all parameters other than the k_+_k_n_ were kept constant. All kinetic parameters obtained are plotted in **Figure 4D**.

