## Supplementary Information for "Aggregation of the Amyloid-β Peptide (Aβ40) within Condensates Generated through Liquid-Liquid Phase Separation"

**Supplementary Video 1:** Time evolution fluorescence microscopy video displaying the Ostwald ripening and coalescence of Aβ40 condensates within a microdroplet. 1:1 ratio of Aβ40:Cla.

**Supplementary Video 2:** Time evolution confocal microscopy video displaying the Ostwald ripening and coalescence of Aβ40 condensates within a microdroplet. 1:1 ratio of Aβ40:Cla.

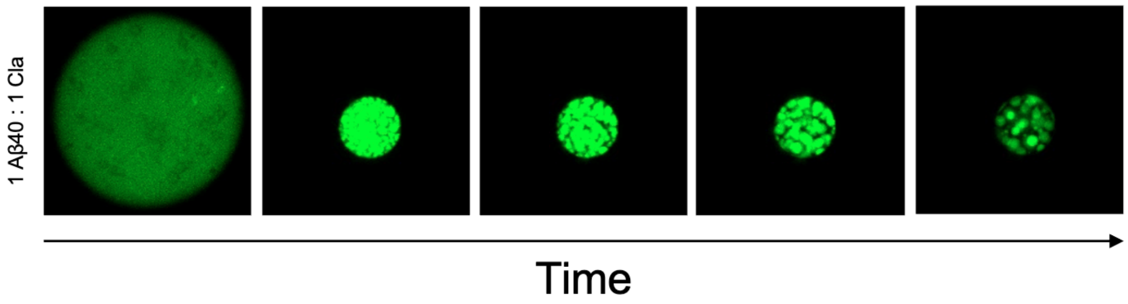


**Supplementary Figure 1. Time evolution of an individual microdroplet displaying the liquid-liquid phase separation of Aβ40.** Images were acquired by confocal microscopy using Alexa Fluor 488 (490 nm excitation wavelength).

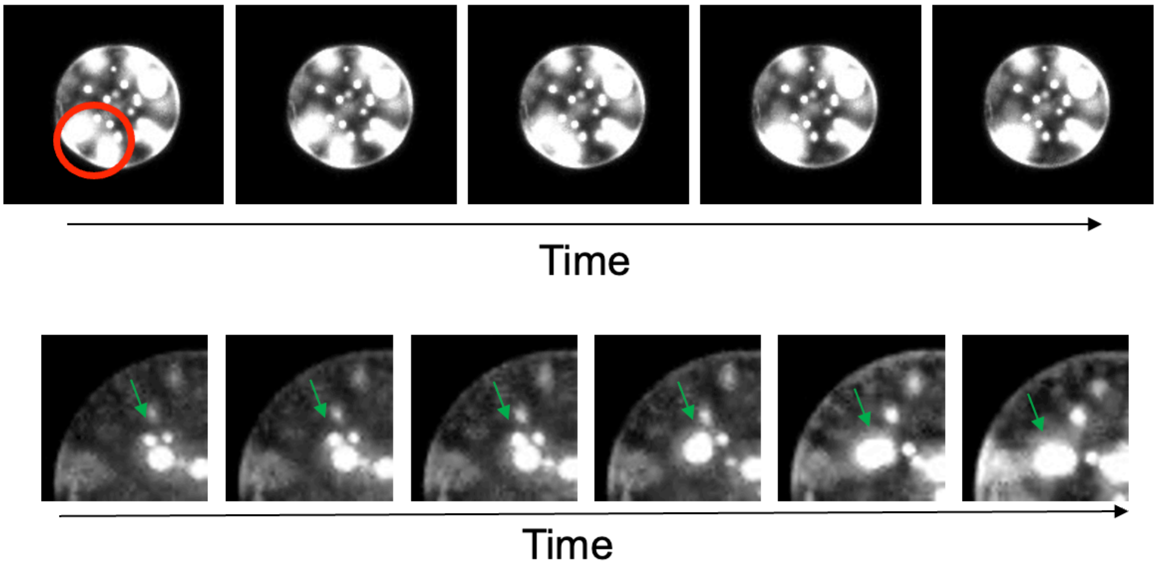


**Supplementary Figure 2. Coalescence of Aβ40 condensates.** Fluorescence imaging of two independent coalescence events occurring between Aβ40 condensates. Images were taken from the time evolution of two independent microdroplets at 1:1 Aβ:claramine stoichiometry.

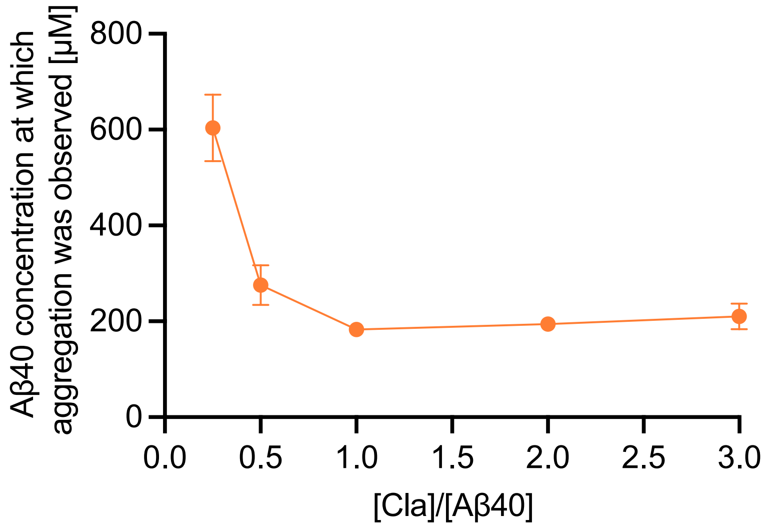


**Supplementary Figure 3. Threshold concentration of Aβ40 required to initiate aggregation within condensates.** Data are shown as mean + SD of n=3.

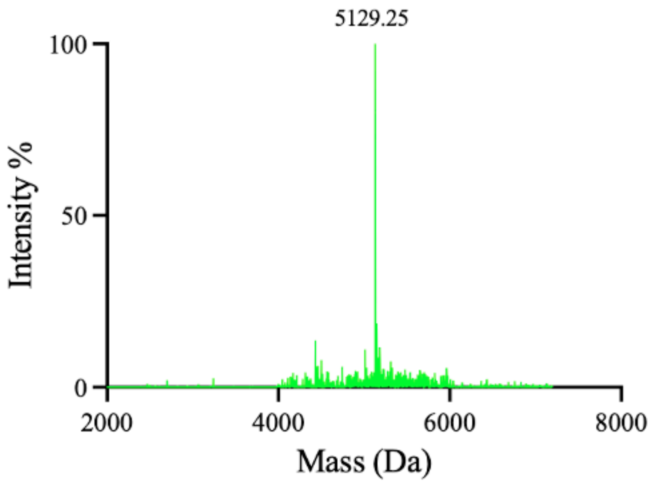


**Supplementary Figure 4. Mass spectrum of Aβ40 conjugated with AlexaFlour 488.**

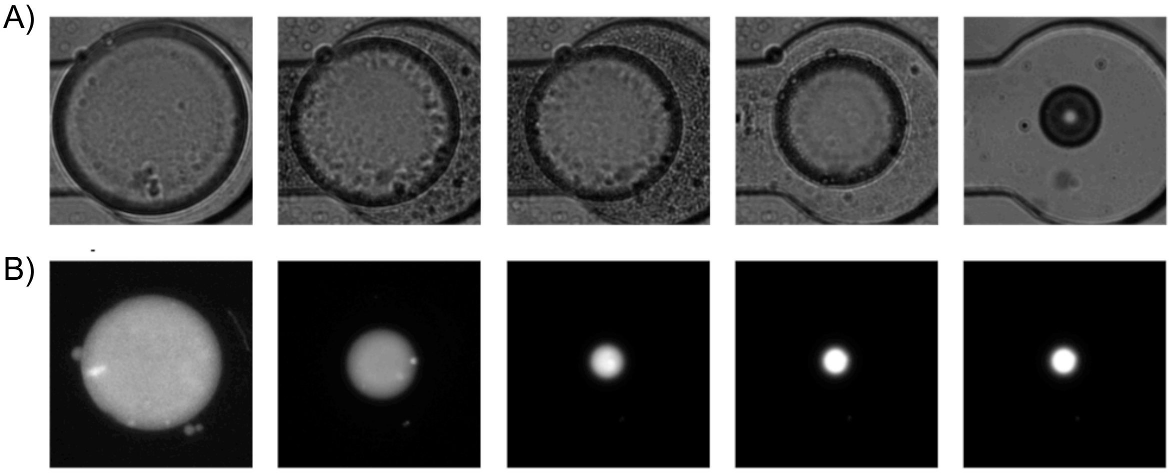


**Supplementary Figure 5. Negative control experiments. (A)** Negative control experiment in which Aβ40 was not present in the sample. The sample contains 7 μM claramine in 50 mM TRIS-HCl, pH 7.4 in the presence of 10% PEG. **(B)** Negative control experiment in which 7 μM Aβ40 was incubated with 7 μM claramine in the absence of 10% PEG.
